## Supplementary figures and tables for "Age and Learning Shapes Sound Representations in Auditory Cortex During Adolescence"

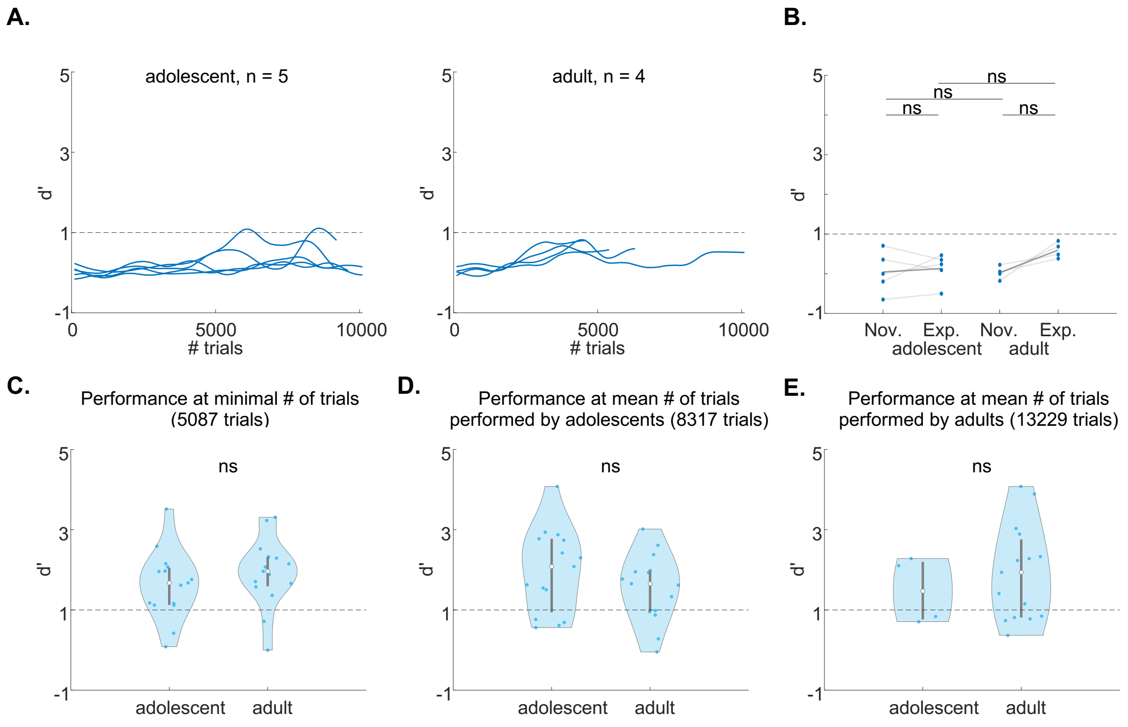


**Supplementary Figure 1-1:** *Behavioral criteria of auditory learning.* **A.** Learning curves per mouse throughout the experiment (n=5 adolescents, n=4 adults). **B.** Discriminability of novice (Nov.; first 100 trials) compared to expert (Exp.; last 100 trials) mice in the adolescent and adult groups (Novice vs Expert — Adolescents, $p = 0.0625$; Adults, $p = 0.0625$, Wilcoxon sign ranked test after Bonferroni correction; Adolescents vs Adults — Novice, $p = 0.4444$, Expert, $p = 0.5476$, Wilcoxon rank sum test after Bonferroni correction). **C.** Discriminability (d') of the easy task at the minimal number of trials (5087 trials) shared between all mice ($z = -1.0370$; $p = 0.2998$, two-sample Wilcoxon rank sum test). ***D.*** Discriminability (d') of the easy task at the mean number of trials of adolescent mice (8317 trials) shared between all mice ($n = 15$ per group; $z = 0.9125$; $p = 0.3615$, two-sample Wilcoxon rank sum test). **E.** Same as A. but for the mean number of trials (13229 trials) of adult mice (adolescent $n = 4$; adult $n = 15$; $z = 0.9125$; $p = 0.5304$, two-sample Wilcoxon rank sum test).

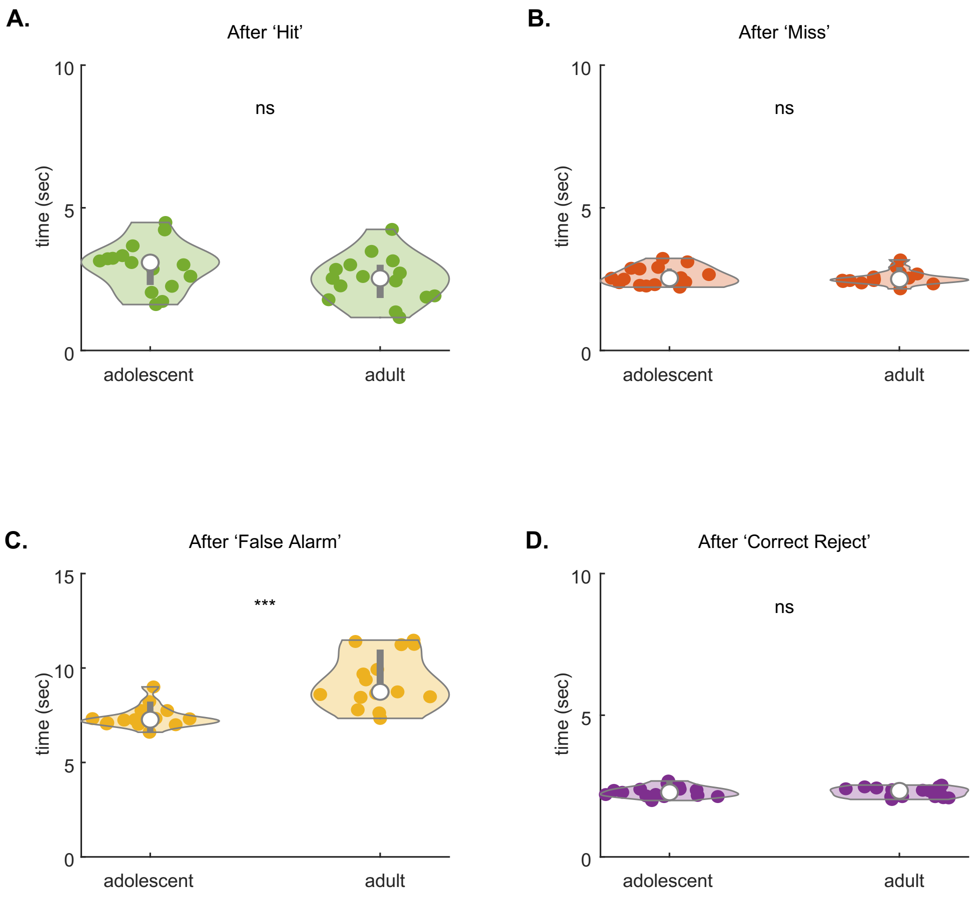


**Supplementary Figure 2-1:** *Inter trial interval after different trial outcomes.* **A.** Average *i*nter trial interval (ITI) per mouse to the next trial after a previous hit ($z = 1.6176$; $p = 0.1057$, two-sample Wilcoxon rank sum test). **B.** Same as A. after a previous miss ($z = 0.0830$; $p = 0.9339$, two-sample Wilcoxon rank sum test). **C.** Same as A. after a previous false alarm hit ($z = -3.9823$; $p = 6.8241e\text{-}05$, two-sample Wilcoxon rank sum test). **D.** Same A. after a previous correct reject hit ($z = 0.3733$; $p = 0.7089$, two-sample Wilcoxon rank sum test).

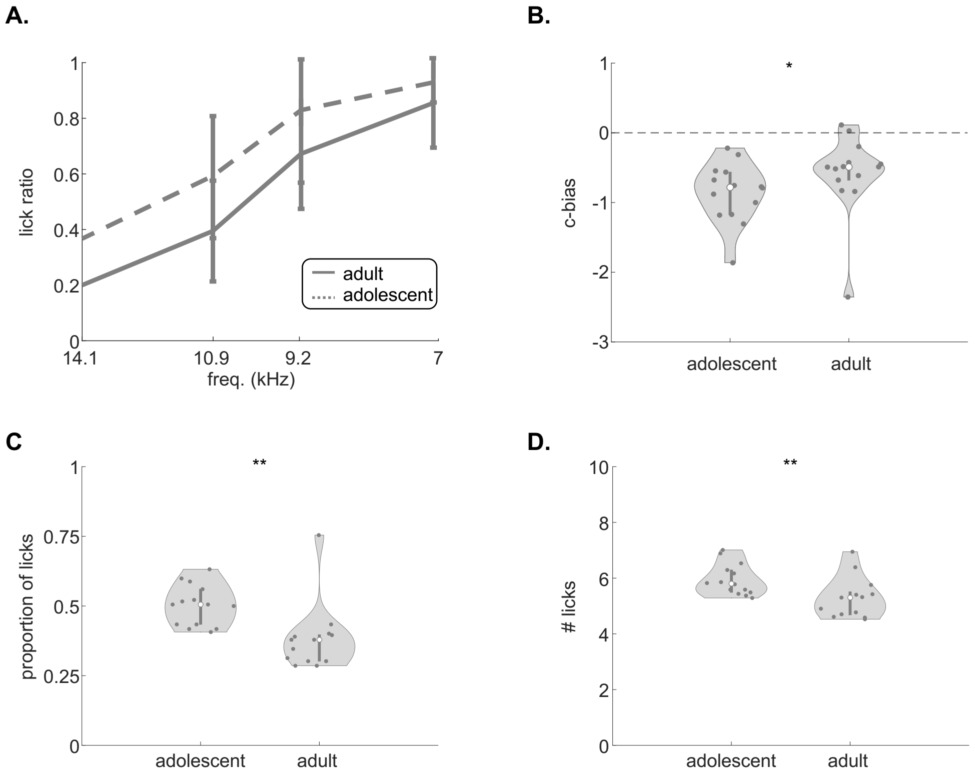


**Supplementary Figure 3-1:** *Lick bias and impulsivity in adolescent and adult mice during head-fixed recordings.* **A.** Average psychometric curve of adult recordings (solid line) and adolescent recordings (dashed line) **B.** Lick bias (i.e., criterion bias) per recording (c-bias: z = -2.1366, p = 0.0326, Wilcoxon rank sum test). **C.** Proportion of licks within ITIs after FAs (z = -2.6447, p = 0.0082, Wilcoxon rank sum test). **D.** Average number of licks during ITIs after FAs (z = -2.7230, p = 0.0063, Wilcoxon rank sum test).

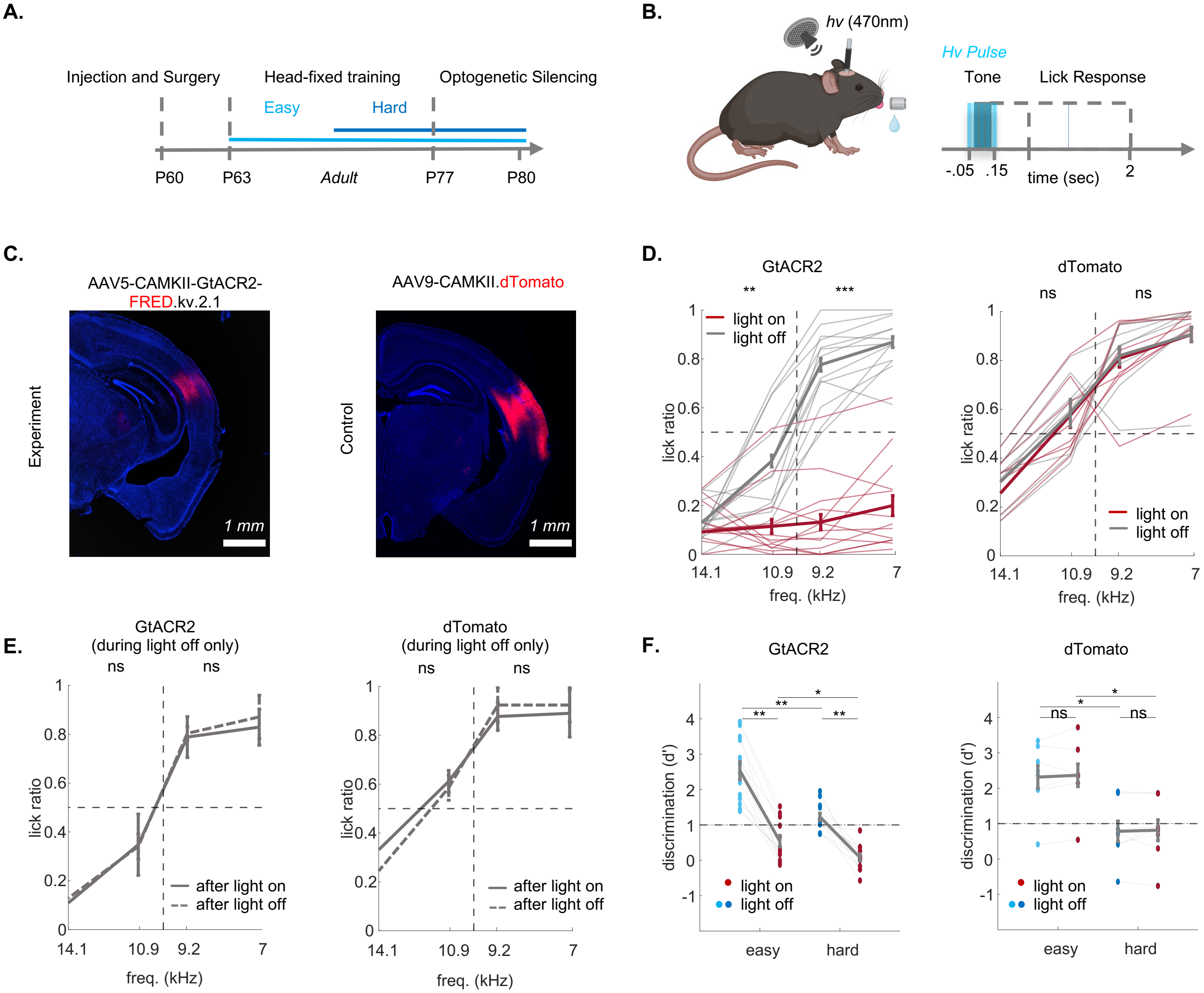


**Supplementary Figure 3-2**: *Auditory Cortex is necessary for task execution in adult mice.* **A.** Experimental design for testing the role of ACx during tone discrimination in expert mice (adults only). **B.** Protocol for transient optogenetic suppression (light pulse duration was -50ms from tone onset, to +50ms from tone offset). **C.** Injection sites and optical fiber implantation for the experimental (GtACR2; top) and control groups (dTomato; bottom). **D.** Lick ratio for Go and No-Go stimuli under light-off conditions (grey) as compared to light-on conditions (red) in experimental (GtACR2, n = 13; left; Go stimuli: p = 0.0001; No-Go stimuli: p = 0.0107; one-sample Wilcoxon sign ranked test after Bonferroni correction) and control mice (dTomato, n = 8; right; Go stimuli: p = 0.4263; No-Go stimuli: p = 0.2953; one-sample Wilcoxon sign ranked test after Bonferroni correction). **E.** Lick ratio under light-off trials after light-on (red) or light off trials (grey) in experimental (GtACR2, n = 13; left; Go stimuli: p = 0.3864; No-Go stimuli: p = 0.2231; one-sample Wilcoxon sign ranked test after Bonferroni correction) and control (dTomato, n = 8; right; Go stimuli: p = 0.58341; No-Go stimuli: p = 0.3214; one-sample Wilcoxon sign ranked test after Bonferroni correction). **F.** Behavioral performance (d') per session under light-off conditions in easy (light blue) and hard (dark blue) task, as compared to light-on conditions (red) in experimental (GtACR2, n = 13; left; easy task: p = 0.0010, hard task: p = 0.0007, two-sample Wilcoxon rank sum test after Bonferroni correction; light-on p = 0.0010; light-off: p = 0.0398, one-sample Wilcoxon sign ranked test after Bonferroni correction) and control mice (dTomato, n = 8; right, easy task: p = 0.9999, hard task: p = 0.7422, two-sample Wilcoxon rank sum test after Bonferroni correction; light on p = 0.0312; light off: p = 0.0234, one-sample Wilcoxon sign ranked test after Bonferroni correction).

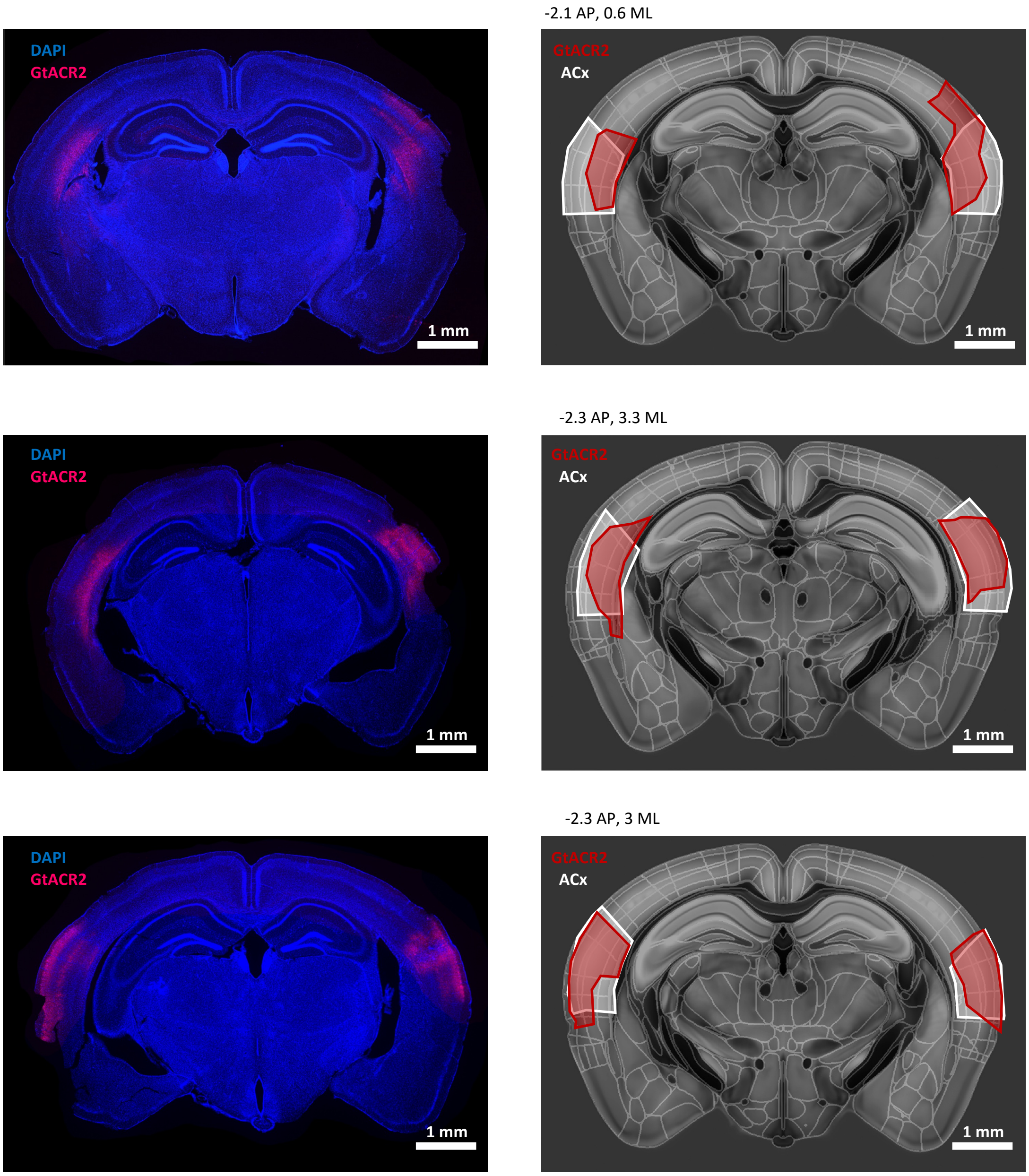


**Supplementary Figure 3-3:** *Verification of GtACR2 expression*. Photomicrograph (left), and reconstruction (right) of the GtACR2 infected area (red) per mouse. Reconstruction followed the coordinates of the Allen-CFF template-atlas. ACx is highlighted in white and GtACR2-expression in red.

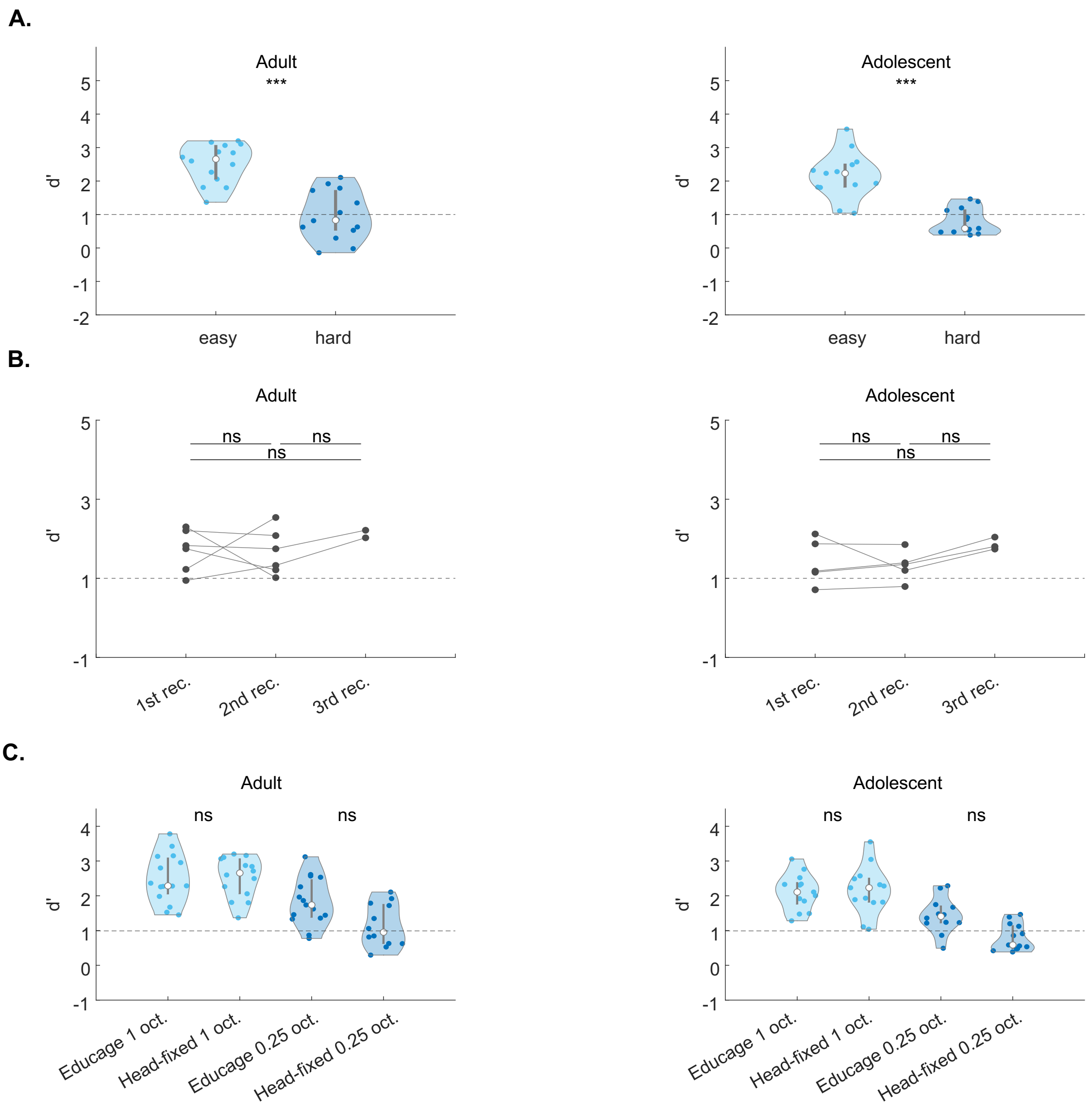


**Supplementary Figure 3-4:** *Adolescent and adult mice performed similarly throughout recordings as well as between the head-fixed configuration and the Educage.* **A.** Behavioral performance (d') in the easy task (light blue) and hard task (dark blue) for adolescent (recording = 13; left) and adult (recording = 14; right) recordings at the behavioral criterion of d' >1 (adolescent mice: signed rank: 91, p = 2.4414e-04; adult mice: signed rank: 105, p = 1.2207e-04, Wilcoxon sign ranked test). Behavioral threshold of d' = 1 highlighted in the dashed line. **B.** Behavioral performance (average d' of the easy and the hard task) for every mouse per recording for adolescent mice (n = 5; left; 1st rec.: p = 0.8125; 2nd rec.: p = 0.9999; 3rd rec.: p = 0.9999, Wilcoxon sign ranked test, after Bonferroni correction), and adult mice (n =6; right; 1st rec.: p = 0. 8438; 2nd rec.: p = 0.9999; 3rd rec.: p = 0.9999, Wilcoxon sign ranked test, after Bonferroni correction). Behavioral threshold of d' = 1 highlighted in the dashed line. **C.** Comparison of behavioral performance in the head-fixed configuration and the Educage for adult mice (left; easy task: p = 0.9960; hard task: p = 0.2159, Kruskal Wallis test after Bonferroni correction), and adolescent mice (right; easy task: p = 0.9973; hard task: p = 0.1505, Kruskal Wallis test after Bonferroni correction) in the easy (light blue) and the hard (dark blue) task.

| Fixed Effects | Estimate | STE | T-Statistic | DF | P-Value | CI lower | CI upper |
|---|---|---|---|---|---|---|---|
| Intercept | 5.441 | 0.2597 | 20.9483 | 1090 | **3.7563e-82** | 4.9314 | 5.9507 |
| Lick Count | -3.4455 | 0.7075 | -4.8697 | 1090 | **1.2827e-06** | -4.8339 | -2.0572 |
| Lick Latency | -0.0086 | 0.0006 | -15.0051 | 1090 | **2.0896e-46** | -0.0098 | -0.0075 |
| d’ | 0.0584 | 0.1102 | 0.5303 | 1090 | 0.5960 | -0.1577 | 0.2746 |
| Count- Latency | 0.0079 | 0.0026 | 3.0258 | 1090 | **0.0076** | 0.0028 | 0.013 |
| Count - d’ | 3.5431 | 1.1445 | 3.0956 | 1090 | **0.006** | 1.2973 | 5.7889 |
| Latency – d’ | -0.0004 | 0.0004 | -0.9635 | 1090 | 0.9999 | -0.0011 | 0.0004 |
| Latency – Count - d’ | -0.0091 | 0.0041 | -2.2524 | 1090 | 0.0735 | -0.0171 | -0.0012 |

**Supplementary Table 1.** *Adolescent and adult mice exhibit different lick behavior in the task.* Linear mixed-effects models of the fixed effects of lick count (until reward or punishment delay), lick latency, cumulative discriminability (d’) (including the interaction effects of lick count and lick latency, lick count and d’, lick latency and d’, and lick latency, lick count and d’) during the minimal number of trials shared between all mice (148 trials; Number of observations = 1098, Fixed effects coefficients = 8, Random effects coefficients = 14, Covariance parameters = 3). Coefficient estimates, STE, T-statistic, degrees of freedom, p-values (adjusted for post-hoc multiple comparisons with Bonferroni-method), lower and higher CI are listed in the table. The model includes random effects coefficients per mouse (11 mice in total) and 3 recordings per mouse (see *methods, equation 8*). Model structure: Lick Count ~ Group * Lick Latency * dprime + (1|Mouse ID) + (1|Recording ID).

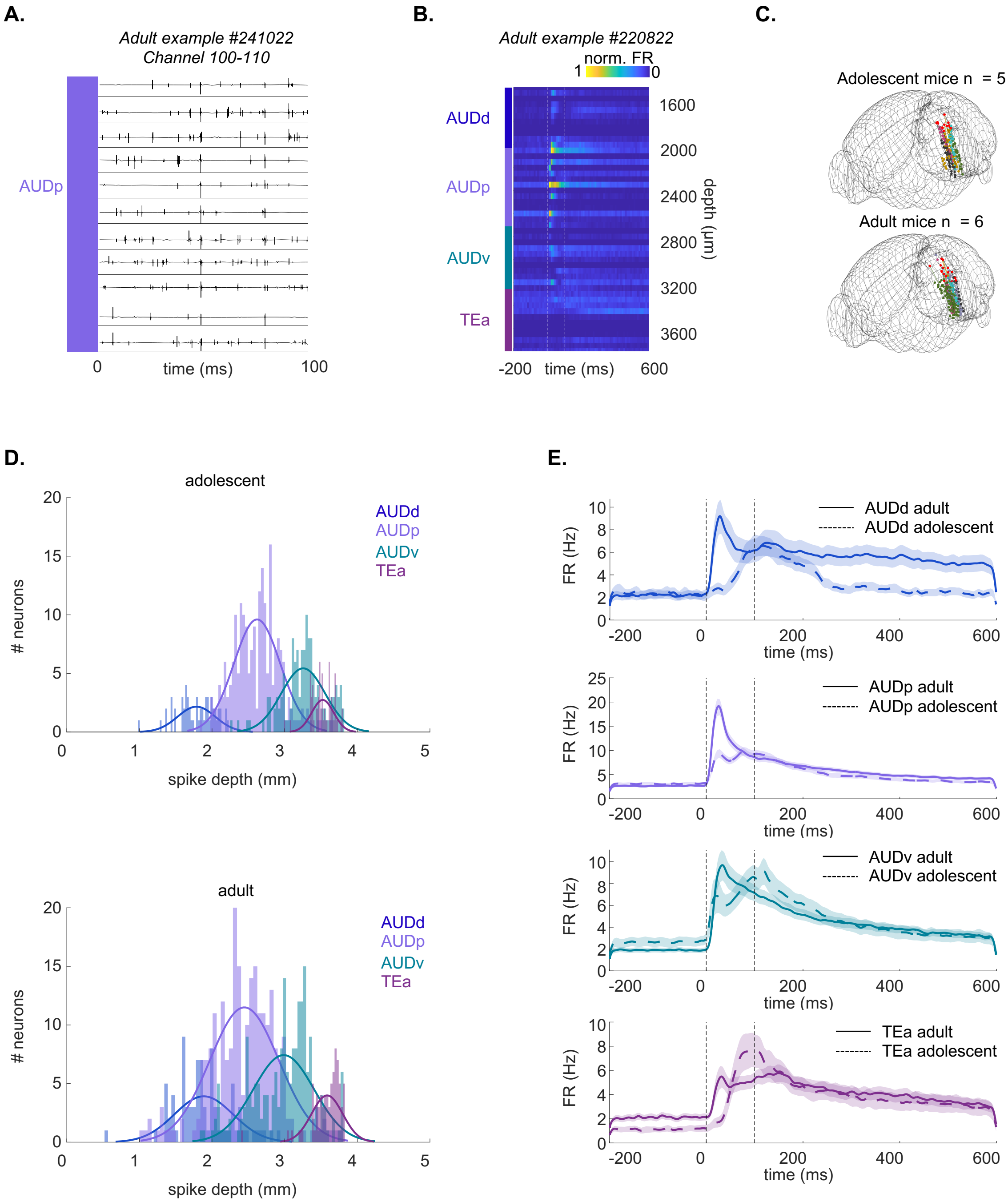


**Supplementary Figure 4-1:** *Probe reconstruction and activity profile across ACx regions.* **A.** Example voltage trace during tone (100ms) across 11 channels. **B.** Example PSTH per depth (50 µm bins) across AUDd, AUDp, AUDv and TEa (3850 maximal depth µm). ***C.*** 3D-Reconstruction of recording sites in adolescent (n = 5; left) and adult (n = 6; right) mice. **D.** Spike-depth of excitatory tone-responsive L5/6 neurons in AUDd, AUDp, AUDv and TEa of adolescent (top) and adult (bottom) recordings. **E.** Population PSTH from – 200ms to 600ms after tone onset in AUDd (blue), AUDp (purple), AUDv (magenta) and TEa (green) for adolescent neurons (dashed line) and adult neurons (solid line).

| Expert | adolescent | | | adult | |
|---|---|---|---|---|---|
| | areas | total | excited | total | excited |
| | AUDd | 177 | 56 (32%) | 154 | 82 (53%) |
| | AUDp | 442 | 210 (48%) | 535 | 245 (46%) |
| | AUDv | 338 | 132 (39%) | 343 | 197 (57%) |
| | TEa | 188 | 65 (35%) | 235 | 75 (32%) |
| **Novice** | **adolescent** | | | **adult** | |
| | areas | total. | excited | total | excited |
| | AUDd | 89 | 22 (25%) | 96 | 17 (18%) |
| | AUDp | 167 | 31 (19%) | 181 | 49 (27%) |
| | AUDv | 302 | 47 (16%) | 180 | 65 (36%) |
| | TEa | 99 | 30 (33%) | 146 | 55 (38%) |

**Supplementary Table 2:** *Neuronal statistics in expert and novice recordings during task engagement.* Acquired single units, acquired tone-excited units (percentage of tone-excited units relative to total units) in the AUDd, AUDp, AUDv, and TEa of adolescent and adult mice in experts (top) and novice (bottom).

| AUDd | adolescent | | adult | | | | | |
| --- | --- | --- | --- | --- | --- | --- | --- | --- |
| neuronal property | Mean | ± STE | Mean | ± STE | Effect size | lower CI | upper CI | p-value |
| spontaneous FR | 2.4274 | 0.4188 | 2.2271 | 0.3668 | 0.1414 | -0.0111 | 0.2997 | 0.9801 |
| evoked FR | 14.5588 | 1.6046 | 22.1731 | 2.9083 | -0.306 | -0.3798 | -0.1396 | 0.1537 |
| FR coeff. var | 2.4484 | 0.2496 | 1.7209 | 0.1238 | 0.4794 | 0.0573 | 0.3294 | **0.0146** |
| latency to peak | 134.5179 | 8.7109 | 132.061 | 16.1264 | 0.1266 | 0.1826 | 0.5423 | **0.0006** |
| FWHM | 275.7321 | 26.1309 | 245.4634 | 24.4123 | 0.1317 | 0.0409 | 0.2443 | 0.2642 |
| min. latency | 73.5638 | 3.3501 | 48.9258 | 3.1867 | 0.7905 | 0.3529 | 0.6238 | **3.6051E-06** |
| % trials resp. | 0.3809 | 0.0364 | 0.4815 | 0.0328 | -0.2993 | -0.3032 | -0.076 | **0.0422** |
| lifetime sparse. | 0.1744 | 0.0291 | 0.1506 | 0.0183 | 0.0401 | -0.3136 | -0.0548 | 0.7897 |
| AUDp | adolescent | | adult | | | | | |
| neuronal property | Mean | ± STE | Mean | ± STE | Effect size | lower CI | upper CI | p-value |
| spontaneous FR | 3.158 | 0.2654 | 2.7504 | 0.2546 | 0.1752 | -0.0103 | 0.2951 | **0.0115** |
| evoked FR | 24.5512 | 1.6812 | 39.4108 | 2.6007 | -0.432 | -0.4069 | -0.1391 | **5.7054E-05** |
| FR coeff. var | 1.6895 | 0.0872 | 1.4396 | 0.084 | 0.2913 | 0.0522 | 0.3326 | **0.0004** |
| latency to peak | 102.981 | 7.9174 | 50.4 | 3.0841 | 0.7259 | 0.1981 | 0.5378 | **1.0347E-11** |
| FWHM | 241.2571 | 13.7176 | 150.2408 | 12.0679 | 0.4569 | 0.0357 | 0.2445 | **8.7880E-10** |
| min. latency | 47.4148 | 1.8017 | 31.4164 | 1.5054 | 0.6936 | 0.3569 | 0.6333 | **5.605E-12** |
| % trials resp. | 0.517 | 0.0208 | 0.6 | 0.0209 | -0.2354 | -0.2853 | -0.0645 | **0.0026** |
| lifetime sparse. | 0.1459 | 0.0115 | 0.1875 | 0.0118 | -0.2739 | -0.3028 | -0.0485 | **0.0049** |
| AUDv | adolescent | | adult | | | | | |
| neuronal property | Mean | ± STE | Mean | ± STE | Effect size | lower CI | upper CI | p-value |
| spontaneous FR | 2.627 | 0.4173 | 1.9547 | 0.2387 | 0.2687 | -0.0121 | 0.2867 | 0.1084 |
| evoked FR | 21.8639 | 2.8512 | 24.4575 | 1.9949 | -0.1752 | -0.3839 | -0.1324 | 0.4306 |
| FR coeff. var | 1.8877 | 0.1089 | 1.7797 | 0.1009 | 0.1175 | 0.0477 | 0.3122 | 0.0927 |
| latency to peak | 126.1818 | 9.6388 | 135.5939 | 10.1844 | -0.0138 | 0.1951 | 0.5405 | 0.2089 |
| FWHM | 226.5985 | 16.5937 | 241.7817 | 15.1074 | -0.0613 | 0.0334 | 0.2419 | 0.7051 |
| min. latency | 59.9072 | 2.3282 | 51.9611 | 2.1164 | 0.2969 | 0.3359 | 0.6227 | **0.0062** |
| % trials resp. | 0.4461 | 0.0253 | 0.5074 | 0.023 | -0.1811 | -0.2962 | -0.0823 | 0.1396 |
| lifetime sparse. | 0.1593 | 0.0153 | 0.2033 | 0.0127 | -0.3125 | -0.3089 | -0.0527 | **0.0038** |
| TEa | adolescent | | adult | | | | | |
| neuronal property | Mean | ± STE | Mean | ± STE | Effect size | lower CI | upper CI | p-value |
| spontaneous FR | 1.2426 | 0.2315 | 2.1943 | 0.2731 | -0.5523 | -0.0096 | 0.303 | **0.0059** |
| evoked FR | 17.567 | 2.5111 | 16.5377 | 1.7923 | -0.0353 | -0.4066 | -0.1551 | 0.8271 |
| FR coeff. var | 1.9063 | 0.1606 | 2.1603 | 0.2278 | -0.0467 | 0.0585 | 0.3122 | 0.9088 |
| latency to peak | 170.0645 | 18.1797 | 145.8133 | 15.4745 | 0.2663 | 0.2071 | 0.5479 | 0.2447 |
| FWHM | 259.6935 | 25.6747 | 297.2267 | 23.7387 | -0.1590 | 0.038 | 0.2512 | 0.4505 |
| min. latency | 73.5202 | 2.8156 | 59.9417 | 2.8259 | 0.5712 | 0.3687 | 0.6166 | **0.0013** |
| % trials resp. | 0.4412 | 0.0375 | 0.4515 | 0.0347 | -0.0491 | -0.3016 | -0.0768 | 0.909 |
| lifetime sparse. | 0.2156 | 0.0238 | 0.1819 | 0.0206 | 0.2167 | -0.3118 | -0.0521 | 0.1658 |

**Supplementary Table 3**: *Adolescent and adult expert mice have distinct firing properties in different sub-regions***.** Mean and standard error (STE), mean effect size (robust Cohen's D), lower and upper Confidence Interval (CI) and p-value (Wilcoxon rank-sum test, adjusted for multiple comparisons with Bonferroni method) of the average baseline FR (Hz), evoked FR (Hz), coefficient of variance of FR, latency to peak of maximal FR (ms), full-width-half maximum of peak FR (ms), minimal latency of first spike (ms), fraction of responsive trials, lifetime sparseness of all adolescent and adult neurons from tone-onset to 50ms after tone offset across all stimuli in AUDd, AUDp, and AUDv, and TEa (significant p-values are highlighted in bold).

| adolescent | P-Value** | | | | | |
|---|---|---|---|---|---|---|
| neuronal property | AUDd - AUDp | AUDd - AUDv | AUDd - TEa | AUDp - AUDv | AUDp - TEa | AUDv - TEa |
| spontaneous FR | 0.3379 | 0.9998 | 0.3018 | **0.0793** | **0.0004** | 0.189 |
| evoked FR | **0.0276** | 0.5874 | 0.9983 | 0.2043 | **0.0367** | 0.6856 |
| FR coeff. var | **0.0078** | 0.3519 | 0.5517 | 0.2063 | 0.3573 | 0.9992 |
| latency to peak | **0.0001** | 0.0878 | 0.9894 | **0.0001** | **0.0001** | 0.1681 |
| FWHM | 0.7022 | 0.4202 | 0.998 | 0.8836 | 0.8076 | 0.5188 |
| min. latency | **0.0001** | **0.0263** | 0.9819 | **0.0002** | **0.0001** | **0.0045** |
| % trials resp. | **0.014** | 0.4845 | 0.6757 | 0.1778 | 0.3250 | 0.9992 |
| lifetime sparse. | 0.9998 | 0.9453 | **0.0255** | 0.8245 | **0.0016** | **0.027** |
| adult | P-Value** | | | | | |
| neuronal property | AUDd - AUDp | AUDd - AUDv | AUDd - TEa | AUDp - AUDv | AUDp - TEa | AUDv - TEa |
| spontaneous FR | 0.999 | 0.4036 | 0.9081 | 0.2066 | 0.7793 | 0.0985 |
| evoked FR | **0.0004** | 0.9843 | 0.6787 | **0.0001** | **0.0001** | 0.3436 |
| FR coeff. var | **0.0119** | 0.9605 | 0.7967 | **0.0038** | **0.0003** | 0.4040 |
| latency to peak | **0.0001** | 0.6264 | 0.0672 | **0.0001** | **0.0001** | 0.3092 |
| FWHM | **0.0023** | 0.9068 | 0.1614 | **0.0001** | **0.0001** | 0.2782 |
| min. latency | **0.0001** | 0.8888 | **0.0409** | **0.0001** | **0.0001** | 0.0744 |
| % trials resp. | **0.0192** | 0.9373 | 0.9163 | **0.0114** | **0.0018** | 0.5453 |
| lifetime sparse. | 0.2722 | **0.0203** | 0.3678 | 0.4004 | 0.9963 | 0.8102 |

** Kruskal Willis Test after Tukey-Kramer correction for multiple comparisons

**Supplementary Table 4:** *Adolescent and adult firing properties of expert mice are distinct between different sub-regions*. P-values of Kruskal Willis Test after Tukey-Kramer correction for multiple comparisons of the average baseline FR (Hz), evoked FR (Hz), coefficient of variance of FR, latency to peak of maximal FR (ms), full-width-half maximum of peak FR (ms), minimal latency of first spike (ms), fraction of responsive trials, lifetime sparseness of all adolescent and adult neurons from tone-onset to 50ms after tone offset across all stimuli AUDd – AUDp, AUDd – AUDv, AUDd – TEa, AUDp – AUDv, AUDp – Tea, and AUDv –Tea (significant p-values are highlighted in bold).

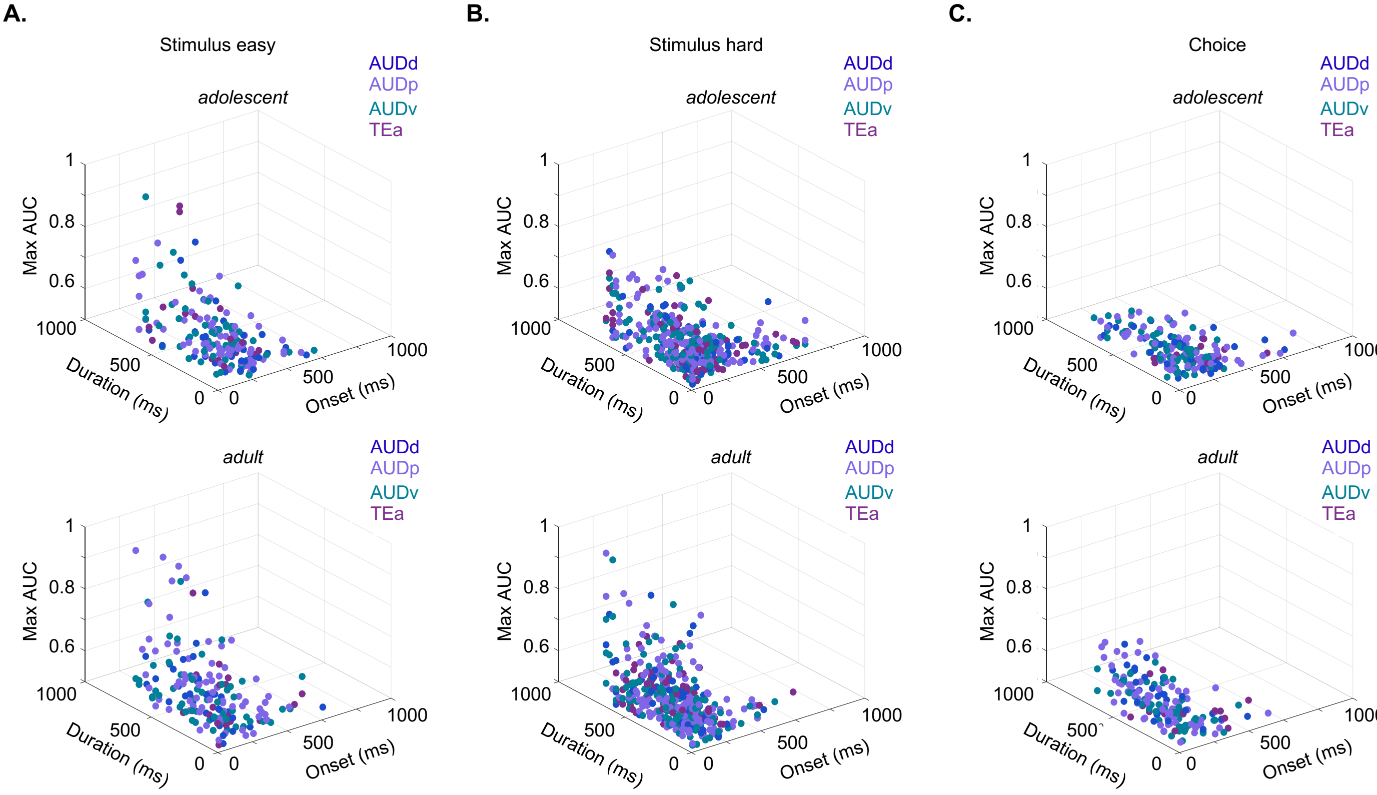


**Supplementary Figure 4-2**. *The neuronal discriminability of stimulus- and choice related activity is similar across auditory sub-regions.* **A.** Onset-latency of discriminability (ms), duration of discriminability (ms), and maximal discriminability (AUC) of neurons that showed significant discriminability (exceeded 3 STD of the shuffled distribution) in the easy task (adolescent neurons = 178 (93%), mice = 4, recordings = 6; adult n = 346 (97%), mice = 4, recordings = 8; adolescent: onset-latency of discriminability: p = 0.5310, duration of discriminability: p = 0.5418, maximal discriminability: p = 0.7212; adult: onset-latency of discriminability: p = 0.3810, duration of discriminability: p = 0.6105, maximal discriminability: p = 0.9115; Friedman test, correct for multiple comparisons). **B**. Same as ‘A’, but in the hard task (adolescent neurons = 399 (93%), mice = 5, recordings = 10; adult n = 544 (97%), mice = 6, recordings = 12; adolescent: onset-latency of discriminability: p = 0.9402, duration of discriminability: p = 0.3388, maximal discriminability: p = 0.6685; adult: onset-latency of discriminability: p = 0.2425, duration of discriminability: p = 0.5700, maximal discriminability: p = 0.1011; Friedman test, correct for multiple comparisons). **C.** Same as ‘A’, but for choice-related activity (adolescent neurons = 181 (95%), mice = 4, recordings = 9; adult n = 339 (95%), mice = 4, recordings = 7; adolescent: onset-latency of discriminability: p = 0.3975, duration of discriminability: p = 0.6823, maximal discriminability: p = 0.2866; adult: onset-latency of discriminability: p = 0.0881, duration of discriminability: p = 0.8185, maximal discriminability: p = 0.2501; Friedman test, correct for multiple comparisons), across the AUDd, AUDp, AUDv, TEa.

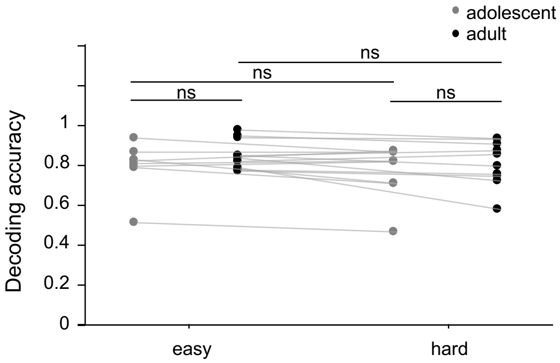


**Supplementary Figure 5-1:** *Decoding accuracy of the first 200ms after the response window.* LDA decoding accuracy the easy and the hard task of adolescent and adult mice 200ms after the response window.

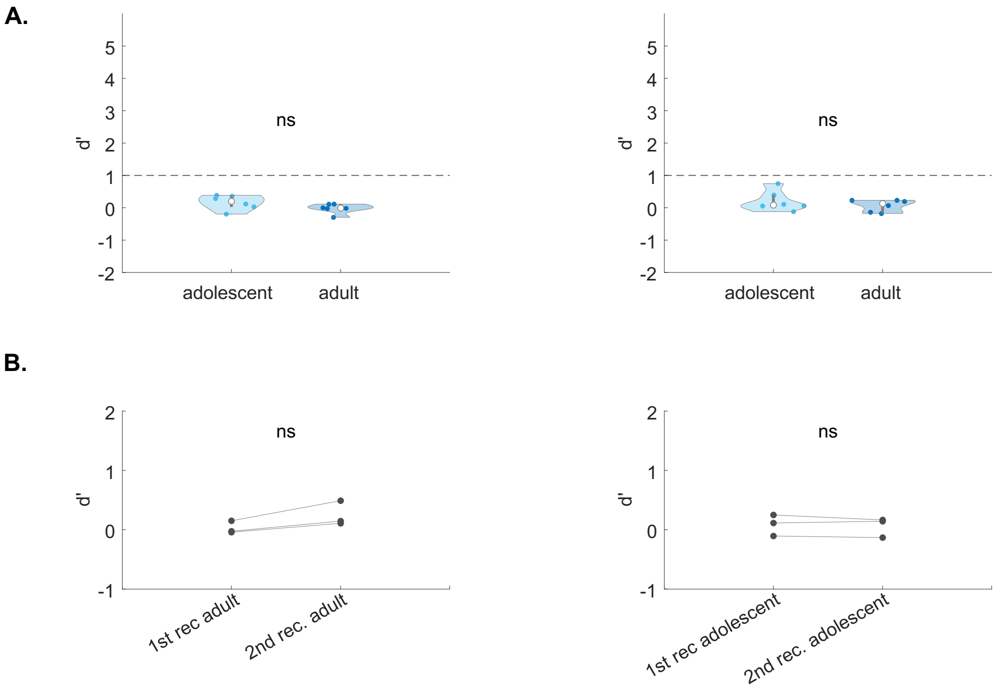


**Supplementary Figure 6-1:** *Behavioral performance of novice mice.* **A.** Behavioral performance (d') in the easy task (light blue) and hard task (dark blue) for adolescent (n = 6; left) and adult (n = 6; right) mice at the behavioral criterion of d' >1 (adolescent mice: signed rank: 19, p = 0.0938; adult mice: signed rank: 15, p = 0.4375, Wilcoxon sign ranked test). Behavioral threshold of d' = 1 highlighted in the dashed line. **B.** Behavioral performance (average d' of the easy and the hard task) for every mouse per recording for adult mice (n = 3; left; signed rank: 0, p = 0.2500 Wilcoxon sign ranked test). adolescent mice (n = 3; right; signed rank: 0, p = 0.2500, Wilcoxon sign ranked test). Behavioral threshold of d' = 1 highlighted in the dashed line.

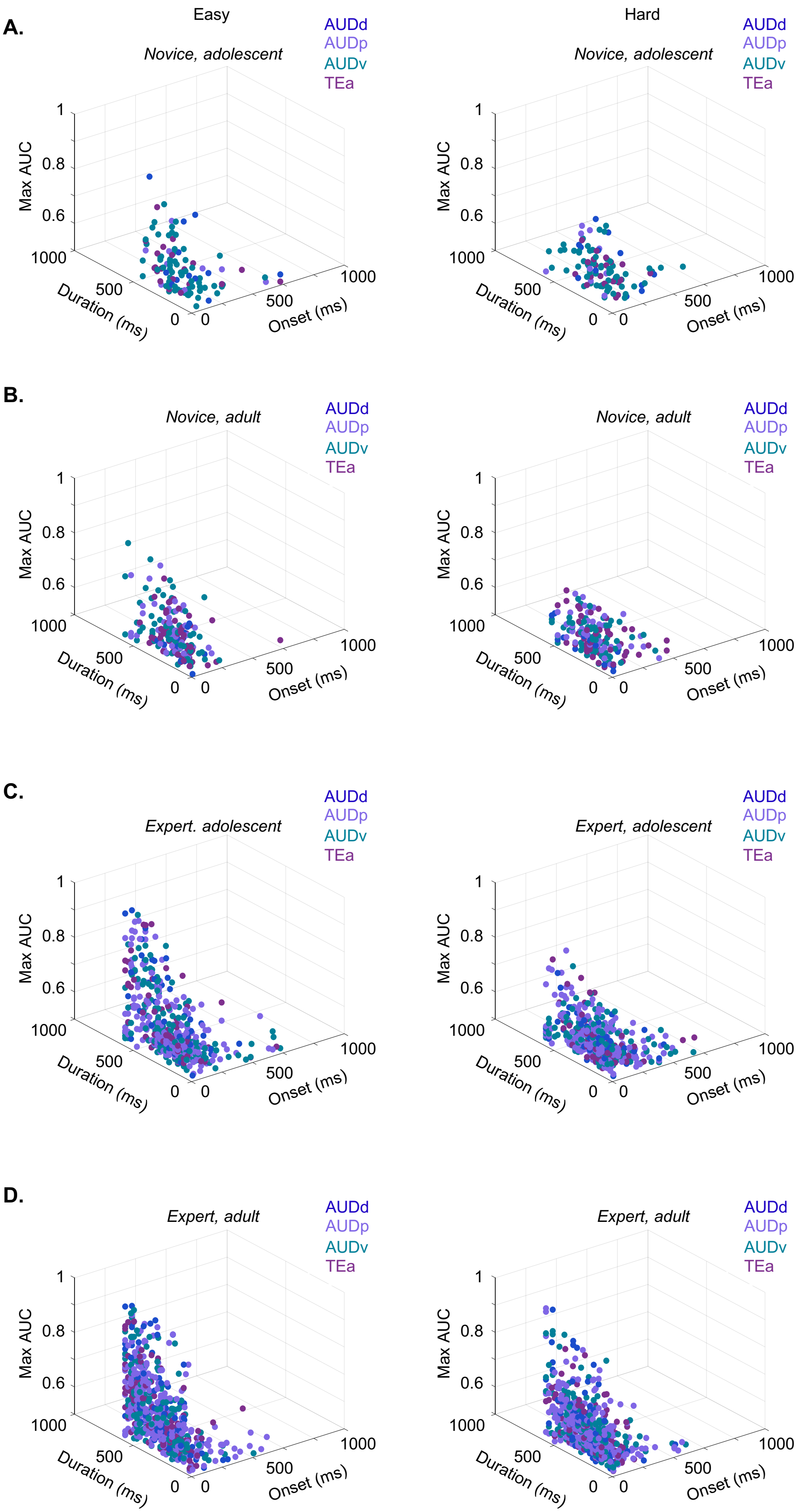
A.
Easy
Hard
Novice, adolescent
Novice, adolescent
AUDd
AUDp
AUDv
TEa
Max AUC
1
0.8
0.6
1000
500
0
Duration (ms)
0
500
1000
Onset (ms)
B.
Novice, adult
Novice, adult
C.
Expert. adolescent
Expert, adolescent
D.
Expert, adult
Expert, adult

**Supplementary Figure 6-2**. *The neuronal discriminability of Easy and Hard Go and No-Go are distributed similar across auditory sub-regions in adolescent and adult novice and expert mice* **A.** Onset-latency of discriminability (ms), duration of discriminability (ms), and maximal discriminability (AUC) of neurons that showed significant discriminability (exceeded 3 STD of the shuffled distribution) in novice, adolescent mice across the AUDd, AUDp, AUDv, TEa. Left: easy task (n = 108 (83%), mice = 3, recording = 6; onset-latency of discriminability: p = 0.2422, duration of discriminability: p =0.5639, maximal discriminability: p = 0.2062; Friedman test, correct for multiple comparisons). Right: hard task (n = 108 (83%), mice = 3, recording = 6, onset-latency of discriminability: p = 0.6294, duration of discriminability: p = 0.0693, maximal discriminability: p = 0.0858; Friedman test, correct for multiple comparisons). **B.** Same as 'A' for novice, adult mice. Left: easy task (n = 179 (97%), mice = 3, recording = 6, onset-latency of discriminability: p = 0.8335, duration of discriminability: p = 0.8013, maximal discriminability: p = 0.1900; Friedman test, correct for multiple comparisons). Right: hard task (n = 181 (96%), mice = 3, recording = 6, onset-latency of discriminability: p = 0.2180, duration of discriminability: p = 0.3388, maximal discriminability: p = 0.0648; Friedman test, correct for multiple comparisons). **C.** Same as 'A' for expert, adolescent mice. Left: easy task (n = 450 (97%), mice = 5, recording = 13, onset-latency of discriminability: p = 0.0918, duration of discriminability: p = 0.4020, maximal discriminability: p = 0.0698; Friedman test, correct for multiple comparisons). Right: hard task (n = 440 (95%), mice = 5, recording = 13, onset-latency of discriminability: p = 0.9911, duration of discriminability: p = 0.9939, maximal discriminability: p = 0.4058; Friedman test, correct for multiple comparisons). **D.** Same as 'A' for expert, adult mice. Left: easy task (n = 598 (99%), mice = 6, recording = 14, onset-latency of discriminability: p = 0.6807, duration of discriminability: p = 0.7223, maximal discriminability: p = 0.7557; Friedman test, correct for multiple comparisons). Right: hard task (n = 589 (98%), mice = 6, recording = 14, onset-latency of discriminability: p = 0.7141, duration of discriminability: p = 0.4084, maximal discriminability: p = 0.6365; Friedman test, correct for multiple comparisons).

| AUDd | adolescent | | adult | | | | | |
|---|---|---|---|---|---|---|---|---|
| neuronal property | Mean | ± STE | Mean | ± STE | Effect size | lower CI | upper CI | p-value |
| spontaneous FR | 2.4274 | 0.4188 | 2.2271 | 0.3668 | 0.3224 | -1.0907 | 1.9771 | 0.9646 |
| evoked FR | 14.5588 | 1.6046 | 22.1731 | 2.9083 | 0.323 | -0.4649 | 1.1014 | 0.4924 |
| FR coeff. var | 2.4484 | 0.2496 | 1.7209 | 0.1238 | -0.2152 | -0.9768 | 0.6759 | 0.3410 |
| latency to peak | 134.5179 | 8.7109 | 132.061 | 16.1264 | -0.0601 | -1.2314 | 0.773 | 0.7230 |
| FWHM | 275.7321 | 26.1309 | 245.4634 | 24.4123 | -0.0333 | -0.7347 | 0.701 | 0.3998 |
| min. latency | 73.5638 | 3.3501 | 48.9258 | 3.1867 | -0.5259 | -1.2905 | 0.1873 | 0.1265 |
| % trials resp. | 0.3809 | 0.0364 | 0.4815 | 0.0328 | 0.1885 | -0.5429 | 1.2357 | 0.7397 |
| lifetime sparse. | 0.1744 | 0.0291 | 0.1506 | 0.0183 | 0.5616 | -0.0983 | 1.7234 | 0.3410 |

| AUDp | adolescent | | adult | | | | | |
|---|---|---|---|---|---|---|---|---|
| neuronal property | Mean | ± STE | Mean | ± STE | Effect size | lower CI | upper CI | p-value |
| spontaneous FR | 3.158 | 0.2654 | 2.7504 | 0.2546 | 0.9234 | 0.5075 | 1.4603 | **1.41592E-05** |
| evoked FR | 24.5512 | 1.6812 | 39.4108 | 2.6007 | 0.8286 | 0.4288 | 1.4088 | **0.0015** |
| FR coeff. var | 1.6895 | 0.0872 | 1.4396 | 0.084 | -0.6202 | -1.0043 | -0.2827 | **0.0012** |
| latency to peak | 102.981 | 7.9174 | 50.4 | 3.0841 | -0.5161 | -1.0001 | -0.0189 | **0.0403** |
| FWHM | 161.2571 | 13.7176 | 150.2408 | 12.0679 | -0.1835 | -0.5979 | 0.393 | **0.0495** |
| min. latency | 47.4148 | 1.8017 | 31.4164 | 1.5054 | -0.4746 | -0.9192 | -0.1092 | **0.0406** |
| % trials resp. | 0.517 | 0.0208 | 0.6 | 0.0209 | 0.667 | 0.2199 | 1.1501 | **0.0007** |
| lifetime sparse. | 0.1459 | 0.0115 | 0.1875 | 0.0118 | 0.3354 | -0.2348 | 0.8616 | 0.1258 |

| AUDv | adolescent | | adult | | | | | |
|---|---|---|---|---|---|---|---|---|
| neuronal property | Mean | ± STE | Mean | ± STE | Effect size | lower CI | upper CI | p-value |
| spontaneous FR | 2.627 | 0.4173 | 1.9547 | 0.2387 | 0.1787 | -0.2081 | 0.6092 | **0.0963** |
| evoked FR | 21.8639 | 2.8512 | 24.4575 | 1.9949 | -0.5624 | -0.929 | -0.2553 | **0.0186** |
| FR coeff. var | 1.8877 | 0.1089 | 1.7797 | 0.1009 | 0.1859 | -0.2146 | 0.6288 | 0.1282 |
| latency to peak | 126.1818 | 9.6388 | 135.5939 | 10.1844 | 0.7395 | 0.3299 | 1.1785 | **0.0001** |
| FWHM | 226.5985 | 16.5937 | 241.7817 | 15.1074 | 0.518 | 0.1243 | 1.1608 | **0.0002** |
| min. latency | 59.9072 | 2.3282 | 51.9611 | 2.1164 | 0.8709 | 0.4572 | 1.4074 | **5.4742E-06** |
| % trials resp. | 0.4461 | 0.0253 | 0.5074 | 0.023 | -0.2375 | -0.5761 | 0.0878 | 0.1676 |
| lifetime sparse. | 0.1593 | 0.0153 | 0.2033 | 0.0127 | 0.3121 | -0.0957 | 0.6979 | **0.0608** |

| TEa | adolescent | | adult | | | | | |
|---|---|---|---|---|---|---|---|---|
| neuronal property | Mean | ± STE | Mean | ± STE | Effect size | lower CI | upper CI | p-value |
| spontaneous FR | 1.2426 | 0.2315 | 2.1943 | 0.2731 | 0.8305 | 0.1538 | 2.1521 | **0.0082** |
| evoked FR | 17.567 | 2.5111 | 16.5377 | 1.7923 | -0.6609 | -0.9696 | -0.2934 | **0.0108** |
| FR coeff. var | 1.9063 | 0.1606 | 2.1603 | 0.2278 | -0.0567 | -0.457 | 0.6441 | 0.8526 |
| latency to peak | 170.0645 | 18.1797 | 145.8133 | 15.4745 | 0.3679 | -0.0907 | 1.0797 | **0.0440** |
| FWHM | 299.6935 | 25.6747 | 277.2267 | 23.7387 | 1.707 | 1.0083 | 3.2231 | **8.99288E-06** |
| min. latency | 73.5202 | 2.8156 | 59.9417 | 2.8259 | 0.5511 | 0.002 | 1.1864 | **0.0112** |
| % trials resp. | 0.4412 | 0.0375 | 0.4515 | 0.0347 | -0.0406 | -0.4834 | 0.4156 | 0.8385 |
| lifetime sparse. | 0.2156 | 0.0238 | 0.1819 | 0.0206 | 0.3157 | -0.3936 | 0.9299 | 0.7058 |

**Supplementary Table 5**: *Novice, adolescent and adult mice have distinct firing properties in different sub-regions***.** Mean and standard error (STE), mean effect size (robust Cohen's D), lower and upper Confidence Interval (CI) and p-value (Wilcoxon rank-sum test, adjusted for multiple comparisons with Bonferroni method) of the average baseline FR (Hz), evoked FR (Hz), coefficient of variance of FR, latency to peak of maximal FR (ms), full-width-half maximum of peak FR (ms), minimal latency of first spike (ms), fraction of responsive trials, lifetime sparseness of all adolescent and adult neurons from tone-onset to 50ms after tone offset across all stimuli in AUDd, AUDp, and AUDv, and TEa (significant p-values are highlighted in bold).

| adolescent | P-Value** | | | | | |
|---|---|---|---|---|---|---|
| neuronal property | AUDd - AUDp | AUDd - AUDv | AUDd - TEa | AUDp - AUDv | AUDp - TEa | AUDv - TEa |
| spontaneous FR | 0.3238 | 0.5109 | 0.1699 | 0.9421 | 0.9336 | 0.6709 |
| evoked FR | 0.6254 | 0.9512 | 0.9985 | 0.6134 | **0.0418** | 0.8233 |
| FR coeff. var | 0.2319 | 0.8251 | 0.3252 | 0.3928 | 0.0998 | 0.5767 |
| latency to peak | 0.188 | 0.8251 | 0.9998 | **0.0002** | **0.0449** | 0.7502 |
| FWHM | 0.7308 | 0.3703 | 0.2985 | **0.0009** | **0.0032** | 0.9723 |
| min. latency | 0.3989 | 0.7007 | 0.7987 | **0.0008** | **0.0137** | 0.9999 |
| % trials resp. | 0.2022 | 0.7372 | 0.2238 | 0.4637 | 0.0996 | 0.5149 |
| lifetime sparse. | 0.9927 | 0.9657 | 0.8218 | 0.9944 | 0.0649 | 0.9267 |

| adult | P-Value** | | | | | |
|---|---|---|---|---|---|---|
| neuronal property | AUDd - AUDp | AUDd - AUDv | AUDd - TEa | AUDp - AUDv | AUDp - TEa | AUDv - TEa |
| spontaneous FR | 0.4679 | 0.9932 | 0.993 | 0.0533 | 0.3267 | 0.8491 |
| evoked FR | 0.9866 | 0.263 | **0.0141** | **0.0096** | **0.0001** | 0.2526 |
| FR coeff. var | 0.9344 | **0.0143** | **0.0047** | **0.0032** | **0.0006** | 0.9345 |
| latency to peak | 0.8167 | 0.1522 | 0.1139 | 0.3118 | **0.0279** | 0.9939 |
| FWHM | 0.3574 | 0.2036 | 0.0662 | 0.9806 | **0.0413** | 0.848 |
| min. latency | 0.16 | **0.0009** | **0.0068** | 0.0884 | **0.0392** | 0.8875 |
| % trials resp. | 0.9998 | **0.0798** | **0.0174** | **0.0021** | **0.0001** | 0.804 |
| lifetime sparse. | 0.9933 | 0.9319 | 0.9764 | 0.9665 | 0.9972 | 0.9932 |

** Kruskal Willis Test after Tukey-Kramer correction for multiple comparisons

**Supplementary Table 6:** *Adolescent and adult firing properties of novice mice are distinct between different sub-regions*. P-values of Kruskal Willis Test after Tukey-Kramer correction for multiple comparisons of the average baseline FR (Hz), evoked FR (Hz), coefficient of variance of FR, latency to peak of maximal FR (ms), full-width-half maximum of peak FR (ms), minimal latency of first spike (ms), fraction of responsive trials, lifetime sparseness of all adolescent and adult neurons from tone-onset to 50ms after tone offset across all stimuli AUDd – AUDp, AUDd – AUDv, AUDd – TEa, AUDp – AUDv, AUDp – Tea, and AUDv –Tea (significant p-values are highlighted in bold).

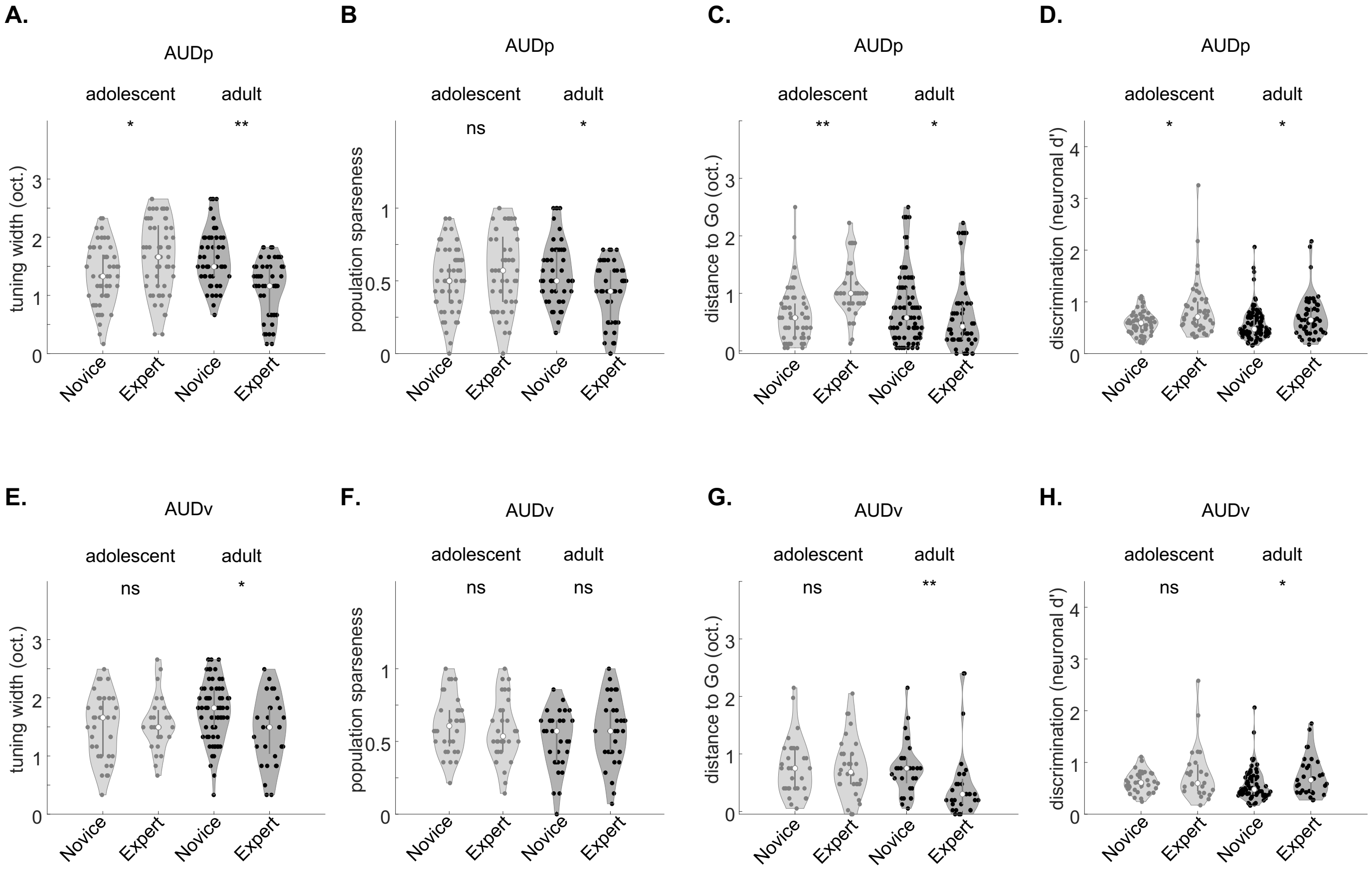


**Supplementary Figure 7-1**: *Learning related changes in neuronal tuning properties in primary and secondary auditory cortex*. **A**. Tuning bandwidth in AUDp at 62 dB SPL of neurons in adolescents and adults. Side by side comparisons of novice versus experts. (adolescents p = 0.0438, adults p = 0.0001, Kruskal Willis Test after Tukey-Kramer correction for multiple comparisons). **B.** Same as 'A' for the population sparseness in AUDp (adolescents p = 0.5724, adults p = 0.0066, Kruskal Willis Test after Tukey-Kramer correction for multiple comparisons). **C.** Same as 'A' for the distance (in octaves) between the best-frequency of each neuron to the easy Go-stimulus in AUDp (adolescents p = 0.0001, adults p = 0.0201, Kruskal Willis Test after Tukey-Kramer correction for multiple comparisons). **D.** Same as 'A' for the average neuronal d' of pairs of frequencies limited to the learned frequency spectrum in AUDp (adolescents p = 0.0471, adults p = 0.0321, Kruskal Willis Test after Tukey-Kramer correction for multiple comparisons). **E-H**. Same as A-D but in the AUDv. **E.** Adolescents p = 0.9982, adults p = 0.0427, Kruskal Willis Test after Tukey-Kramer correction for multiple comparisons. **F**. Adolescents p = 0.8841, adults p = 0.8031, Kruskal Willis Test after Tukey-Kramer correction for multiple comparisons. **G.** Adolescents p = 0.9910, adults p = 0.0042, Kruskal Willis Test after Tukey-Kramer correction for multiple comparisons. **H.** Adolescents p = 0.9438, adults p = 0.0108, Kruskal Willis Test after Tukey-Kramer correction for multiple comparisons.

| Group | | | | | |
| --- | --- | --- | --- | --- | --- |
| Expert | | adolescent | | adult | |
| | areas | total | excited | total | excited |
| | AUDd | 29 | 1 (0.03%) | 64 | 13 (0.2%) |
| | AUDp | 147 | 44 (0.30%) | 111 | 27 (0.24%) |
| | AUDv | 137 | 35 (0.26%) | 109 | 25 (0.23%) |
| | TEa | 35 | 0 (0%) | 124 | 19 (0.15%) |
| Novice | | adolescent | | adult | |
| | areas | total. | Excited | total | Excited |
| | AUDd | 89 | 6 (0.07%) | 96 | 8 (0.08%) |
| | AUDp | 167 | 25 (0.15%) | 181 | 24 (0.13%) |
| | AUDv | 302 | 44 (0.15%) | 180 | 45 (0.25%) |
| | TEa | 99 | 17 (0.17%) | 146 | 46 (0.32%) |

**Supplementary Table 7:** *Neuronal statistics during passive FRA protocol in expert and novice mice.* Acquired single units, acquired tone-modulated units, and percentage of modulated units to all acquired units in the AUDd, AUDp, AUDv, and TEa of adolescent and adult mice during passive-listening recordings.

| AUDp | adolescent | | adult | | | | | |
| --- | --- | --- | --- | --- | --- | --- | --- | --- |
| neuronal property | Mean | ± STE | Mean | ± STE | Effect size | lower CI | upper CI | p-value |
| spontaneous FR | 15.0108 | 2.552 | 16.1657 | 2.0109 | -0.1182 | -0.6373 | 0.3627 | 0.8636 |
| evoked FR | 33.9654 | 8.0969 | 40.81 | 4.012 | -0.0807 | -0.5791 | 0.4458 | 0.7709 |
| FR coeff. var | 0.847 | 0.1456 | 0.6278 | 0.1104 | 0.5127 | 0.0149 | 1.0775 | 0.0646 |
| latency to peak | 110.8182 | 8.3515 | 61.963 | 17.0012 | 0.4648 | 0.0048 | 0.9211 | **0.0472** |
| FWHM | 178.3409 | 21.0212 | 68.5185 | 26.5098 | 0.6418 | 0.2081 | 1.0021 | **0.0032** |
| min. latency | 55.5352 | 6.6073 | 48.058 | 4.6893 | 0.242 | -0.2302 | 0.7889 | **0.0389** |
| % trials resp. | 0.6335 | 0.0827 | 0.5972 | 0.5320 | 0.0716 | -0.473 | 0.507 | 0.9951 |
| lifetime sparse. | 0.4198 | 0.0458 | 0.3383 | 0.3580 | 0.2868 | -0.2248 | 0.7884 | 0.1048 |
| AUDv | adolescent | | adult | | | | | |
| neuronal property | Mean | ± STE | Mean | ± STE | Effect size | lower CI | upper CI | p-value |
| spontaneous FR | 18.9295 | 1.8448 | 8.7466 | 3.8163 | 0.47 | -0.0115 | 0.8816 | **0.0174** |
| evoked FR | 46.1598 | 5.3813 | 21.9135 | 8.8535 | 0.6166 | 0.0396 | 1.2171 | **0.0068** |
| FR coeff. var | 1.0532 | 0.232 | 0.9158 | 0.1627 | 0.2143 | -0.3916 | 0.9365 | 0.1785 |
| latency to peak | 109.4857 | 19.6319 | 89.68 | 12.9514 | 0.3197 | -0.3407 | 0.8796 | 0.18106 |
| FWHM | 146.0857 | 30.3333 | 119.2 | 26.5399 | 0.1272 | -0.4042 | 0.7038 | **0.0439** |
| min. latency | 60.4915 | 8.3884 | 49.5169 | 4.3829 | 0.2985 | -0.2216 | 0.8729 | **0.0433** |
| % trials resp. | 0.6946 | 0.0804 | 0.3925 | 0.0545 | 0.7253 | 0.1905 | 1.5257 | **0.0027** |
| lifetime sparse. | 0.3773 | 0.0533 | 0.2565 | 0.0367 | 0.4362 | -0.0431 | 1.1493 | **0.0451** |

**Supplementary Table 8**: *Adolescent and adult mice have distinct firing properties across different sub-regions of ACx—passive listening***.** Mean and standard error, mean effect size (robust Cohen's D), lower and upper Confidence Interval (CI) and p-value (Wilcoxon rank-sum test) of the average baseline FR (Hz) (AUDp vs. AUDv: adolescent p = 0.8551; adult p = 0.9711), evoked FR (Hz) (AUDp vs. AUDv: adolescent p = 0.4125; adult p = 0.9954), coefficient of variance of FR (AUDp vs. AUDv: adolescent p = 0.4354; adult p = 0.8800), latency to peak of maximal FR (ms) (AUDp vs. AUDv: adolescent p = 0.5871; adult p = 0.9985), full-width-half maximum of peak FR (ms) (AUDp vs. AUDv: adolescent p = 0.7223; adult p = 0.4628), minimal latency of first spike (ms) (AUDp vs. AUDv: adolescent p = 0.5936; adult p = 0.5669), fraction of responsive trials (AUDp vs. AUDv: adolescent p = 0.3838; adult p = 0.9924), lifetime sparseness (AUDp vs. AUDv: adolescent p = 0.3792; adult p = 0.9341) of all adolescent and adult neurons from tone-onset to 50ms after tone offset across all stimuli in AUDp, and AUDv (significant p-values are highlighted in bold).

| Group | Expert | | | Novice | | |
|---|---|---|---|---|---|---|
| | Mice | Recording | Neurons | Mice | Recording | Neurons |
| Figure 1 | 15 (adolescent)<br>15 (adult) | none | none | none | none | none |
| Figure 2 | 15 (adolescent)<br>15 (adult) | none | none | none | none | none |
| Figure 3 | 5 (adolescent)<br>6 (adult) | none | none | none | none | none |
| Figure S3-2 | 4 (GtACR2)<br>3 (dTomato) | none | none | none | none | none |
| Figure 4 | 5 (adolescent)<br>6 (adult) | 13 (adolescent)<br>14 (adult) | 1145 (adolescent)<br>1267 (adult) | none | none | none |
| Figure 5 | 5 (adolescent)<br>6 (adult) | 13 (adolescent)<br>14 (adult) | 1145 (adolescent)<br>1267 (adult | none | none | none |
| Figure 6 | 5 (adolescent)<br>6 (adult) | 13 (adolescent)<br>14 (adult) | 1145 (adolescent)<br>1267 (adult) | 3 (adolescent)<br>3 (adult) | 6 (adolescent)<br>6 (adult) | 657 (adolescent)<br>603 (adult) |
| Figure 7 | 4 (adolescent)<br>4 (adult) | 4 (adolescent)<br>4 (adult) | 348 (adolescent)<br>408 (adult) | 3 (adolescent)<br>3 (adult) | 6 (adolescent)<br>6 (adult) | 557 (adolescent)<br>503 (adult) |

**Supplementary Table 9:** *Overview of datasets analyzed.* Number mice, recordings, and neurons per learning stage (left: expert, right: novice) for every figure.
